# Plant–soil feedbacks between arbuscular- and ecto-mycorrhizal communities

**DOI:** 10.1101/228387

**Authors:** Kohmei Kadowaki, Satoshi Yamamoto, Hirotoshi Sato, Akifumi S. Tanabe, Amane Hidaka, Hirokazu Toju

**Author notes:** equal contribution (Kohmei Kadowaki, Satoshi Yamamoto). **Corresponding author:** Kohmei Kadowaki, Center for Ecological Research, Kyoto, University, Otsu, Japan.

## Abstract

Soil microbiomes of adult trees exert species-specific effects on the survival and growth of seedlings^1-6^, yet empirical evidence that such plant–soil microbiome interaction drives seedling community assembly remains scarce. Here we show that mycorrhizal fungal communities determine seedling community assembly by controlling how resident plant communities alter the growth of newly established seedlings. We reciprocally introduced seedling communities of arbuscular-and ecto-mycorrhizal plant species to replicated mesocosms to follow the effects of mycorrhizal type match/mismatch with resident plant communities on seedling growth rates. The growth rates of recruited seedlings were generally higher under resident trees of the same mycorrhizal types than under those of different mycorrhizal types, generating positive plant–soil feedbacks through mycorrhizal-type matching. Such positive effects of matching were linked with seedlings’ greater acquisition rates of mycorrhizal symbionts from matched resident plants than from mismatched plants, and such linkage was pronounced for ecto-mycorrhizal plant species. In contrast, under the condition of mycorrhizal-type matching between resident plants and seedlings (i.e., within-mycorrhizal-type comparison), plant–soil feedback effects varied considerably in their sign and strength among resident–seedling species combinations. Consequently, the assembly of a temperate tree seedling community is driven by a combination of species-specific plant–soil feedbacks and the match/mismatch of mycorrhizal type between resident plants and seedlings.

---

Feedback between plant community assembly and soil biota (or plant–soil feedback) is critical to understanding the dynamics of forest ecosystems such as coexistence and succession^1,2^. Plant–soil feedbacks occur when the effects of soil biota that reside in association with a given plant species are expressed more strongly on conspecific than on heterospecific seedlings^2^. Although many experimental studies have shown that soil microbiota can exhibit such species-specific influences on recruited seedling survival and growth^3-6^, those analyses were restricted to the effects of one plant species on another (i.e., plant-species pairwise interactions). Therefore, we still have limited knowledge of how plant–soil feedback influences seedling performance under mixed plant-species conditions. Furthermore, when plant–soil feedback effects on seedlings have been quantified in greenhouse or in the field, the dynamics of the soil microbiomes have rarely been followed. Consequently, few studies have considered how soil microbiomes affect plant–soil feedbacks through connecting neighboring resident trees and seedlings via networks of interactions^5,7^. Although most previous studies using a single resident species have shown the importance of species-specific feedback effects, the feedback effects under mixed plant-species conditions may have profoundly different effects on seedling community assembly from those predicted under single resident species conditions. Incorporating these realistic properties of plant–microbiome interactions remains a major challenge for understanding plant–soil feedbacks at the community level.

While many experimental studies have shown variability in the sign and strength of plant–soil feedbacks, the results indicate that negative feedbacks are generally limited to arbuscular mycorrhizal (AM) plant species and positive feedbacks are typically observed in ectomycorrhizal (EcM) plant species^7^. The negative feedbacks of AM plant species increase the biomass of fungal species that make the soil less suitable for conspecific seedlings relative to heterospecifics, thereby promoting the coexistence of different AM plant species at the community level^8-10^. In contrast, the positive feedbacks of EcM plant species increase the biomass of fungal species that favor conspecific seedlings over heterospecifics, thereby promoting the dominance of the EcM species within a community^11,12^.

Given that AM and EcM plants co-occur across broad climatic ranges and that EcM forests frequently have an AM understory in temperate zones^12^, matching/mismatching of mycorrhizal type between resident plants and recruited seedlings is of particular importance in considering plant community dynamics^13,14^. Specifically, EcM seedlings may grow faster than AM seedlings when colonizing EcM resident habitats in a forest, generating positive feedbacks as a consequence of mycorrhizal type matching. While many studies have shown variability in the strength and sign of plant–soil feedbacks within the same mycorrhizal type (i.e., AM seedling/AM resident, or EcM seedling/EcM resident conditions), few have quantified plant–soil feedbacks under the situation in which resident plants and colonizing seedlings have mismatched mycorrhizal types^13^. Furthermore, such plant–soil feedbacks might be more profoundly affected by mycorrhizal-type match/mismatch at the community level (i.e., mixed plant-species condition) than at the species level (i.e., single resident species), because the roles of belowground plant–fungal dynamics may be more pronounced at the community level: in mixed plant conditions, multiple seedlings and residents collectively form common mycelial networks belowground through mycorrhizal fungi^15-17^. Previous studies of AM and EcM seedling species implicate that the two plant groups could drive distinct plant–fungal dynamics, in which belowground fungal community structure changes in response to plant community dynamics and where these changes differentially feed back on plant community processes. Considering plant–soil feedbacks within and across mycorrhizal-types and the underlying belowground plant–fungal dynamics will lead to an improved understanding of the consequences of plant–soil feedbacks for plant community assembly.

To determine whether plant–soil feedbacks within and across mycorrhizal types affect seedling community assembly, we experimentally assembled plant–fungal communities, varying mycorrhizal type (AM or EcM) of resident plant communities and colonizing seedling communities in a fully crossed factorial design (Fig. 1). In experimental mesocosms simulating forest stands, we established resident plant communities carrying mycorrhizal inocula (the conditioning phase) and then introduced uninoculated seedling communities into the mesocosms to follow the subsequent growth of the seedlings (the feedback phase). By assessing the feedback effects based on seedling growth patterns within and across mycorrhizal types, we show that the assembly of a temperate tree community is determined by a combination of species-specific plant–soil feedbacks and the match/mismatch of mycorrhizal type between resident plants and seedlings. This study links the observed feedback effects within and across mycorrhizal types to the dynamics of belowground fungal communities, providing a basis for understanding how multiple types of mycorrhizal symbioses drive plant community assembly.

**Figure 1.**
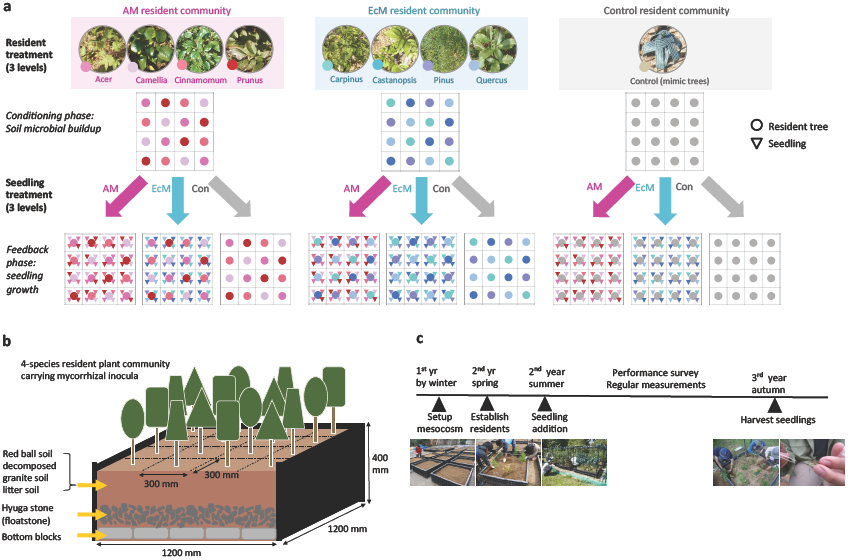
Schematic depiction of the experimental design, mesocosm layout, and the timeline of research. (a) A two-way factorial design of reciprocal invasion experiment, using three by three resident–seedling treatment combinations. The experiment was undertaken following the two time steps: the conditioning phase and the feedback phase. In the conditioning phase, resident communities carrying mycorrhizal inocula were established into the mesocosms filled by soil, each community consisting of 16 resident trees (represented by 4 species × 4 individuals per species, each species filled by different color). In the feedback phase, four seedlings of different species (triangles) were planted at four inter-cardinal positions in each grid cell (*N*= 16 seedlings per species) and were allowed to grow in interaction with resident trees and their belowground fungal associates. For the no-resident treatments, mimic trees made of pole and cheesecloth were planted, and for the control seedling treatments, no seedlings were added to the mesocosms. (b) Illustration of a replicate mesocosm (1.2 m × 1.2 m × 0.4 m height) that consists of multiple soil horizons: artificial bedrock, float-stone, and a standardized mixture of red-ball soil, decomposed granite soil, and litter soil. (c) Timeline represents times when major activities in the study occurred. Photos by authors.

## Results

### Plant–soil feedbacks within and across mycorrhizal types

In the community-scale experiment crossing three resident levels (a four-species AM community, a four-species EcM community, and control [mimic trees]) and three seedling levels (a four-species AM community, a four-species EcM community, and control [no seedlings]) (9 combinations × 4 replicates = 36 mesocosms), we assessed seedling growth patterns to estimate feedbacks effects over two growing seasons (Fig. 1; see *Full methods* in the Supporting Information). For each mesocosm, we averaged seedling species performance by all possible seedling and resident species combinations at the grid level (Fig.1a). The resultant population averages for different seedling species showed that most seedling species exhibited the highest growth in the no-resident control (mimic trees), possibly due to competition for soil nutrients with resident plant community (see *Evidence for resident-seedling competition* and Fig. S1 in the Supporting Information). As such, we defined plant–soil feedback effect as the growth difference of seedlings when grown with conspecific versus heterospecific resident trees (within the same grid cell, Fig. 1a), divided by the growth in the no-resident control (standardizing among different seedling species the potential effects of competition that could affect interspecific comparison of the feedback effects). This index measures the relative advantage of seedlings growing under conspecific vs heterospecific resident species, with the identity of heterospecific resident being used as a contrast varying from heterospecifis of the same mycorrhizal type to those of different mycorrhizal type (see *Feedback effects on seedling growth* in Methods). Based on this formulation, the sign of feedback effect is positive (or negative) when seedlings perform better (or worse) under conspecific relative to heterospecific resident species. Such feedback effects could arise when the effects of soil fungal communities that reside in association with a given plant species would be expressed more strongly on conspecific seedlings than on heterospecific seedlings.

By examining the feedback effects on seedlings for different seedling species when resident communities carried matching versus mismatching inocula, we found that mycorrhizal type matching was a significant predictor of the feedback effects for two AM and three EcM seedling species we examined. When feedback effects were evaluated across matching mycorrhizal types (i.e., AM resident/EcM seedling or EcM resident/AM seedling conditions), the two AM seedling species *Acer* and *Prunus* showed exhibited higher growth in the presence of conspecific relative to heterospecific EcM resident species, indicating positive feedbacks when EcM resident plant species were used as contrasts (i.e., mismatching mycorrhizal type) (Fig. 2). Furthermore, three EcM species *Carpinus*, *Castanopsis* and *Pinus* generally showed positive feedbacks when mismatching resident species (i.e., AM resident species) were used as contrasts for evaluating feedback effects (Fig. 2).

**Figure 2.**
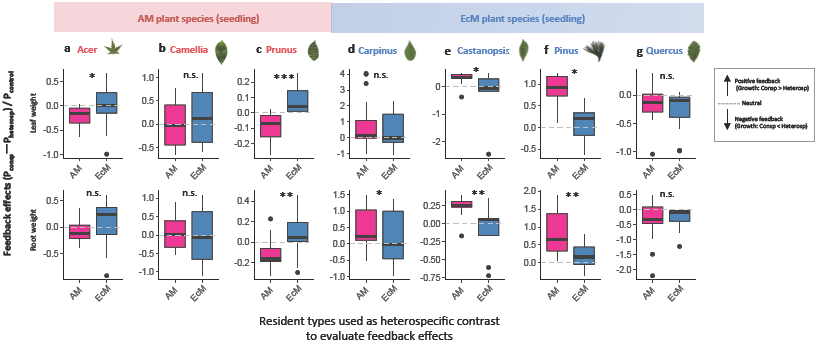
plant–soil feedbacks effects across mycorrhizal types at the community-level. Feedback effect was calculated as the relative increase in seedling performance under conspecific resident species using the formulation: the growth difference when grown under conspecific vs heterospecific resident trees (as contrast) P_consp_–P_heterosp_, divided by the growth under no-resident control treatments P_control_ For each of the three AM seedling species (a-c) ^†^ and four EcM seedling species (d-g), differences in feedback effects were tested across matching vs mismacthing mycorrhizal types (*P < 0.05, ** P < 0.01, *** p < 0.001), using two growth components (dry leaf or root weight). ^†^Note: The results of AM seedling species *Celtis* was not shown as feedback effects were not calculated; this species occurred only as seedling species, but not as resident species; hence, calculation of the feedback effects require a full set of seedling growths measurements under conspecific, heterospecific and control no-resident conditions for the seedling species.

In contrast, when feedback effects were evaluated within the same mycorrhizal type (i.e., AM resident/AM seedling or EcM resident/EcM seedling conditions), AM species often exhibited lower growth in the presence of conspecific relative to heterospecific AM resident species, indicating negative feedbacks when feedback effects were evaluated within AM resident community (Fig. 3). Three EcM species *Carpinus*, *Castanopsis* and *Pinus* showed large variation in the sign and magnitude of feedback effects, depending on resident–seedling species combinations (Fig. 3). One EcM species *Quercus* frequently showed negative feedbacks irrespective of the mycorrhizal types of the resident plants used as heterospecific contrast.

**Figure 3.**
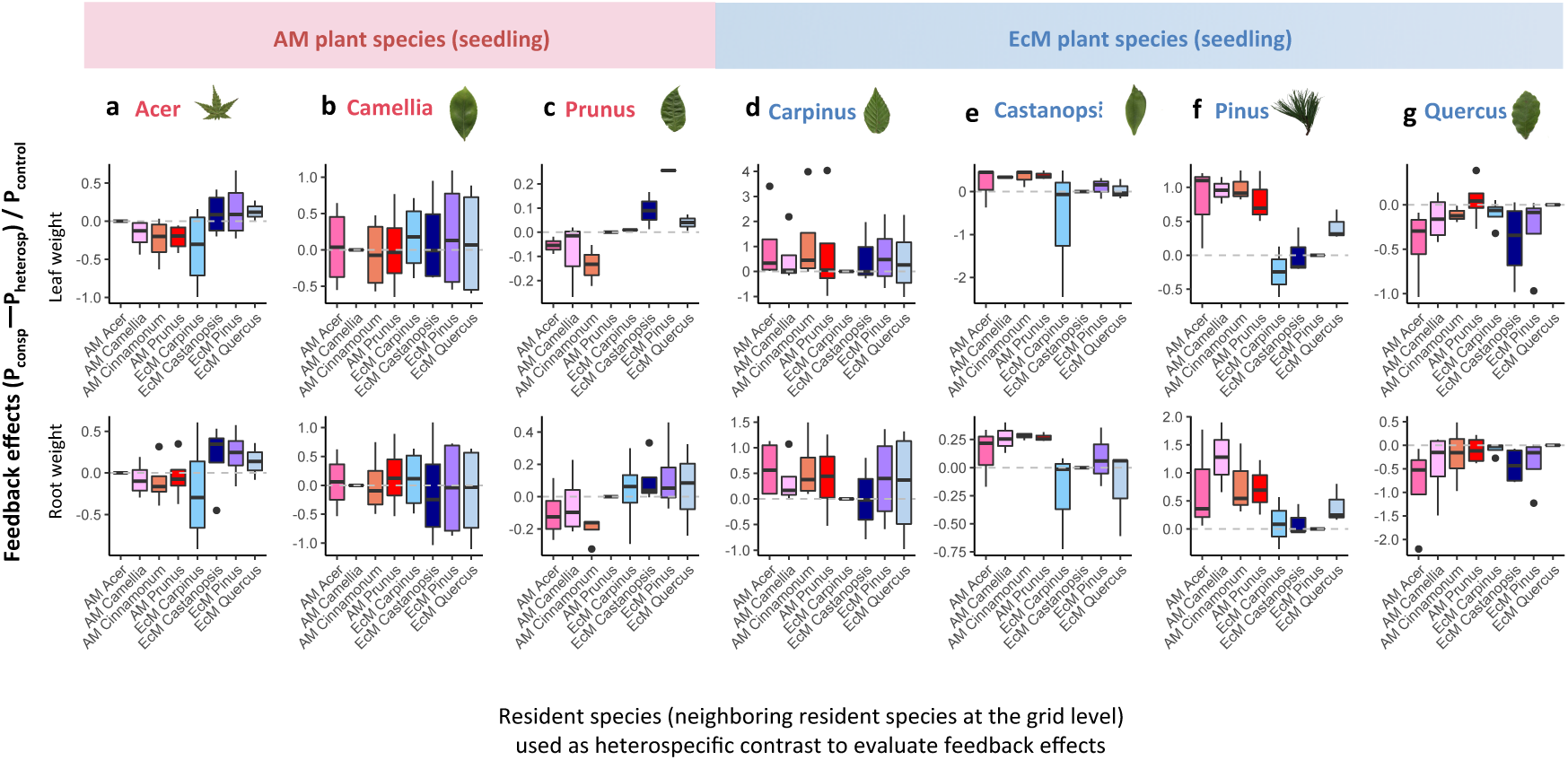
Plant–soil feedbacks effects at the species level. For each of the three AM seedling species (a-c) and four EcM seedling species (d-g), feedback effects were plotted for conditions where different resident species was used as target species (i.e., contrast to seedling growth under conspecific resident species) for the evaluation of feedback effects. The same dataset as Fig.2 was used to illustrate species-pairwise feedback effects.

As a whole, the result shows that mycorrhizal type match was beneficial for both AM and EcM seedling species, and the feedback effects were generally positive when seedling growth rates were compared within matching mycorrhizal types. In contrast, when the feedback effects were compared within the same mycorrhizal types, AM seedling species generally showed negative feedbacks but EcM seedling species showed large variation in the sign and strength of feedbacks (ranging from negative, neutral, to positive feedback effects) depending on the resident–seedling species pair (Fig. S2).

To test for the generality of the effects of mycorrhizal type match/mismatch on the plant–soil feedbacks, we used four different seedling fitness components for calculation of feedback effects, and then analyzed data using linear mixed model with feedback effects as response variable and match/mismatched resident type as a predictor. When all the seedling species were analyzed simultaneously, we confirmed that the results were qualitatively similar among different seedling fitness components, and the effect of mycorrhizal type match/mismatch was robust in all the models of growth components fitted (Table S2). The results show that mycorrhizal-type match might have a strong potential to generate positive feedbacks.

### Sharing of mycorrhizal fungi

To link the observed positive matching effects on feedbacks with increased seedling access to mycorrhizal fungi, we examined whether seedling species had a greater chance of being colonized by shared fungal species with matching resident species (AM resident/AM seedling and EcM resident/EcM seedling treatments) than with mismatching resident species (AM resident/EcM seedling and EcM resident/AM seedling treatments). We found that EcM seedling species shared higher proportions of fungal operational taxonomic units (OTUs; 30.7% based on Morisita-Horn index) with EcM resident species than with AM resident species (11.3%; Fig. 4). In contrast, AM seedling species did not show any preferential association with either resident type, with approximately 14.4–16.3% of OTUs being shared (Fig. 4). This asymmetry resulted in a significant seedling × resident interaction effect on a seedling’s symbiont acquisition rate (Table S3).

**Figure 4.**
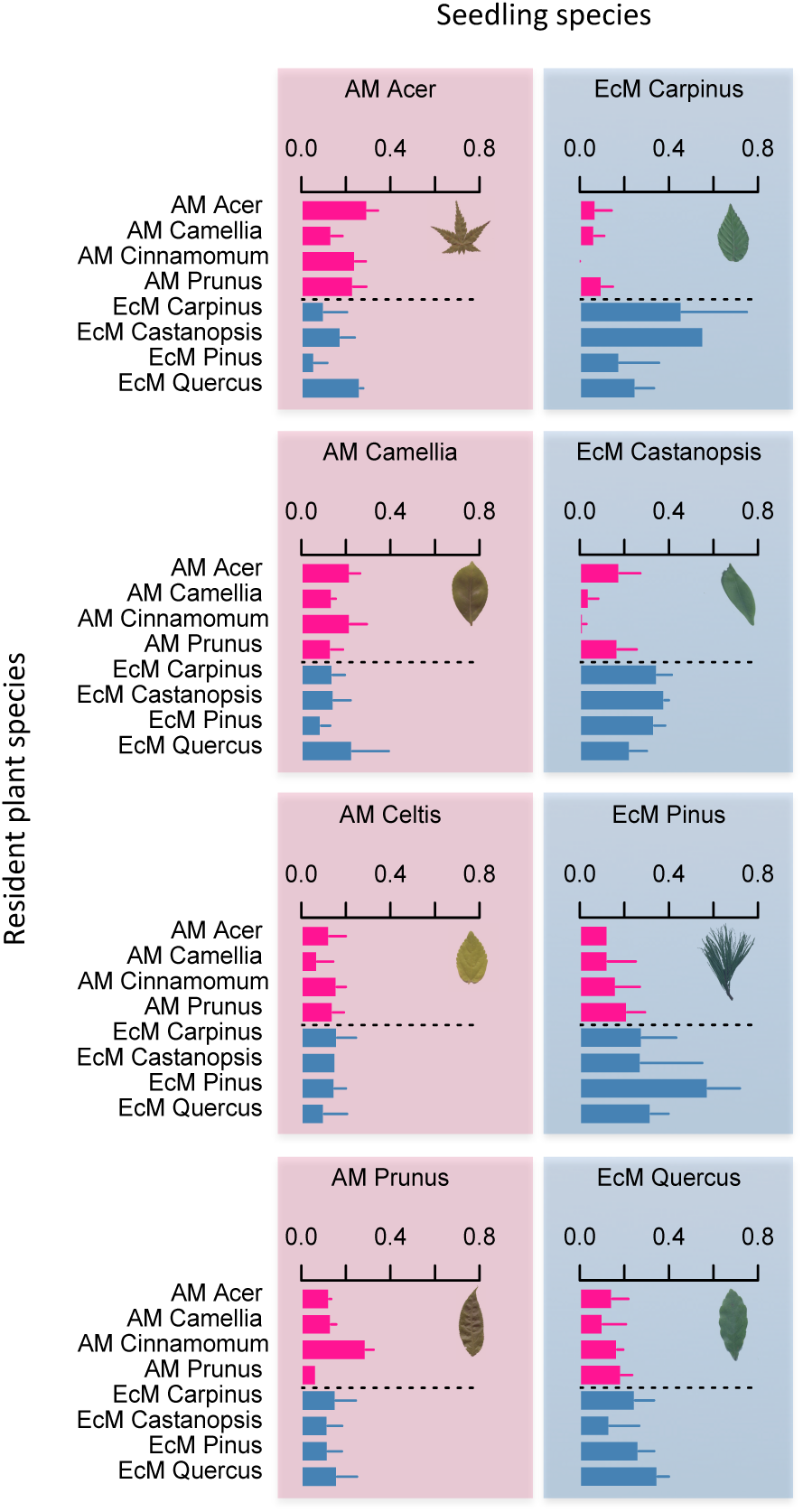
Seedling species’ symbiont acquisition rates. For each resident–seedling species pair in every mesocosm, the proportions of fungal operational taxonomic units (OTUs) of seedling species shared with neighboring different resident species at the time of harvest (mean + 1SE) was evaluated based on 1 – *C*_MH_ (Morisita-Horn similarity).

### Spatial structuring of plant–fungus associations

If fungal community composition becomes similar between resident trees and seedlings at the grid level, one would expect to detect spatial patterns in belowground plant–fungus associations at the mesocosm scale. In our experiment, we used a grid system in each mesocosm to represent all possible species-pairwise resident–seedling interactions (within grid cells) by standardizing the spatial configurations of resident and seedling species (Fig. 1). We took advantage of this grid design to examine how spatial patterns in belowground plant–fungus associations differ depending on resident–seedling treatment combinations (that is, mycorrhizal type match/mismatch).

EcM seedlings were more likely to share fungal OTUs with close resident trees than with distant ones in mesocosms only in the presence of matching EcM resident communities (Fig. 5; Spearman’s rank correlation coefficient (ρ) between fungal community similarity and resident–seedling spatial distance: matching EcM vs. mismatching AM = 0.217 vs. 0.033, *t* = 6.064, df = 4.792, *P* = 0.002, Welch’s two-sample *t*-test). In contrast, AM seedlings showed consistently weaker spatial structuring (Spearman’s ρ: matching AM vs. mismatching EcM = 0.115 vs. 0.090), with no differences depending on resident treatments (*t* = 0.386, df = 2.305, *P* = 0.732). Under EcM resident/EcM seedling conditions, fungal communities were spatially autocorrelated within the distance of 5–50 cm (beyond neighboring grid cells) in mesocosms (results not shown). Under AM resident/AM seedling conditions, however, even nearby grid cells differed in fungal composition, leaving no spatial signal of below-ground plant–fungal associations.

**Figure 5.**
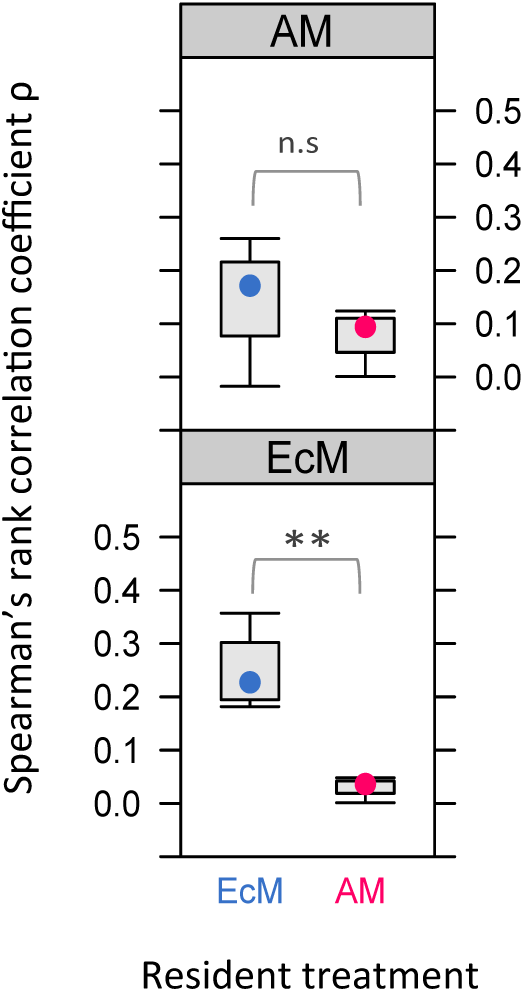
Boxplots of spatial structuring in belowground fungal networks connecting resident and seedling communities. For each resident seedling treatment combination, Spearman’s rank correlation coefficient ρ (ranging from –1 to +1) was estimated between (i) resident-seedling fungal community dissimilarity and (ii) physical distance (separation) within a mesocosm. The upper panel represents AM seedling treatment and the lower panel represents EcM seedling treatment. A high correlation coefficient indicates that close resident–seedling pairs within mesocosms tend to host more similar fungal community composition.

## Discussion

Work to date has shown that soil microbiomes of adult trees exert both species-specific positive and negative effects on the survival and growth of seedlings, but it remains unclear how such feedbacks with mycorrhizal fungal communities shape plant community assembly in mixed plant conditions. Our community-scale experiment demonstrates that the assembly of a temperate tree seedling community is determined by a combination of species-specific plant–soil feedbacks and the match/mismatch of mycorrhizal type between resident plants and seedlings. The results are summarized into three key points. First, resident–seedling mycorrhizal type match/mismatch was a significant predictor of plant–soil feedbacks, benefitting the performance of recruited seedlings when their mycorrhizal type (i.e., AM or EcM) matched those of seedlings (Fig. 2, Table S2). Second, the examined plant species varied in the sign and strength of feedbacks when their seedling performance was evaluated *within* the AM or EcM plant community, suggesting a relatively minor role of species-specific feedbacks within the same type (Fig.3, Fig. S2). Third, mycorrhizal match/mismatch affected the rate of fungal symbiont acquisition and spatial structuring of fungal communities (Figs. 4, 5), which might be linked with the observed feedback effects on seedling community assembly, at least for EcM plant species.

Previous studies have generally reported negative feedbacks in AM symbioses and positive feedbacks in EcM symbioses^4,6,10,11,18,19^. Because those studies focused on feedbacks among different species within a single mycorrhizal plant guild (i.e., AM seedling vs. AM resident or EcM seedling vs. EcM resident conditions), few studies have quantified plant–soil feedbacks in situations in which resident plants and colonizing seedlings are incongruent in terms of their mycorrhizal types (i.e., AM resident/EcM seedling or EcM resident/AM seedling conditions). Bennett *et al*.^13^ used soil inoculum from matching versus mismatching plant species to detect any effect of mycorrhizal type matching and concluded that species-specific feedback effects could play a more important role than mycorrhizal type match/mismatch. They found that AM plant species consistently exhibited negative feedbacks, whereas EcM plant species consistently showed positive feedbacks.

Here we found that, in mixed plant conditions, the sign and strength of feedbacks depend critically on whether we evaluate seedling growth within or across mycorrhizal types. When plant–soil feedbacks are evaluated *within* an AM or EcM plant community, the feedback effects on recruited seedlings varied greatly in the sign of feedbacks depending on resident–seedling species combinations. In contrast, when the feedback effects were compared between mycorrhizal type matching versus mismatching conditions, the sign of plant–soil feedbacks was mostly consistent among different seedling species. The results suggest that the information of mycorrhizal type match/mismatch, along with that of species-specific feedbacks, help us predict seedling community assembly.

Mycorrhizal type matching with the resident plant community was beneficial to colonizing seedlings for both AM and EcM symbioses (Fig. 2). Such beneficial matching effects may have resulted from four non–mutually exclusive mechanisms: (i) more spatially extended and temporally prolonged infection of seedling roots by matching fungal communities than by mismatching ones; (ii) detrimental effects of incompatible mycorrhizal fungi for seedlings that mismatched their mycorrhizal type with resident community; (iii) access to a larger soil nutrient pool made available by compatible fungal networks than by incompatible networks^20^; and (iv) more active transport of nutrients from resource-rich regions of mycelial networks to resource-poor areas^21^ via more structured networks in matching symbioses. The analyses of fungal communities within the mesocosms (Figs. 4, 5) suggested that AM and EcM symbioses were differentiated in their associations at the examined spatial scale, consistent with all these possibilities. We were able to link the dynamics of belowground fungal communities with the observed positive matching effects for EcM seedling species, but not for AM seedling species. Although mechanisms underlying the link between fungal symbiont acquisition and plant–soil feedbacks deserve further investigation, our results highlight the contrasting functional roles of AM versus EcM associations as a predictor of plant community assembly.

Our approach to determining the roles of plant–soil feedbacks is different from previous studies in several ways. First, many previous studies either focused on the effects of live soil inocula associated with a single resident plant species on conspecific or heterospecific seedlings (i.e., plant competition-free conditions)^9,13,18^ or on the effects of one resident plant species on another^16,21-24^. We examined the hypothesis that tree seedlings could exhibit feedbacks at the community level, focusing on resident–seedling interactions across and within mycorrhizal types. By comparing how matching versus mismatching conditions underlie plant–soil feedbacks in mixed plant conditions, our results emphasize that evaluating plant–soil feedbacks both within and across mycorrhizal types will improve our ability to predict seedling community assembly.

Second, most previous studies have used long-lived, large trees as resident species and introduced seedlings underneath^11-13,16^, and such approaches have often left the surrounding vegetation and the soil microbial environment uncontrolled. As a result, the standard way to evaluate plant–soil feedbacks has been the use of fungicide application^6^. This removes the potential effects of soil pathogens, but the treatment was confounded with possible reductions of soil mycorrhizal fungi, which could also act as key agents of feedback effects^15-17^. This study assembled plant–soil communities (i) on identical substrate of a set area, (ii) naturally and *in situ* at a field site, and (iii) for a known period of time to address plant–soil feedbacks and the dynamics of belowground fungal communities in mixed plant conditions (see ref.^25-26^ for an exception). Our study is the first to link plant–soil feedbacks with analysis of belowground plant-fungus associations at the whole ecosystem level and the consequences of the linkages for seedling community assembly.

It is necessary to note, however, some caveats of our study: we did not control for the effects of many other potentially important factors, such as plant species diversity and identity. For the sake of experimental feasibility and tractability, we chose two specific tree communities from each type to test for within/between type feedback effects. If we are to confirm the generality of the findings, further studies should be undertaken to assess how variable these results could be for different sets of AM and EcM species used to build plant–soil communities. Researchers have shown, for example, that plant–soil feedbacks not only affect plant community structure^1,6,8^ (but see ref^23^.) but also are affected by plant species diversity^19,27^. Because many potential mechanisms influencing the functioning of mycorrhizal symbioses (e.g., light intensity, soil fertility ^20,28-30^) must differ between previous pot experiments and our mixed-plant mesocosm experiments, the results of within-type feedbacks should be interpreted with caution.

Understanding whether seedling community assembly is determined by species-specific plant–soil feedbacks or by mycorrhizal type match/mismtatch is critical to predicting plant community assembly and succession^1,30^. Although the idea that mycorrhizal type is a significant predictor of plant community succession is not new^7,31^, it has not previously been tested experimentally at the ecosystem level. Our study presented a first step towards contrasting AM and EcM plant–soil feedbacks in a multi-species context and highlighted the importance of simultaneously examining AM and EcM plant communities. To develop a more comprehensive understanding of plant community assembly, future studies need to quantitatively evaluate the roles of both AM and EcM fungi^31-33^ as well as the diversity and biomass of mycorrhizal, endophytic, and pathogenic fungi in plant root systems^2,34^ in association with various abiotic factors. Incorporating such complexities of real belowground plant–size of the seedlings of the eight speciesfungus interactions will provide a basis for predicting plant community dynamics.

## Methods

### Model system

Using plant species common to warm-temperate forests in Japan, we studied two types of model communities: a four-species AM plant community (*Acer palmatum* Thunb., *Camellia japonica* L., *Cinnamomum camphora* [L.] J. Presl, and *Prunus jamasakura* [Siebold ex Koidz.] H. Ohba) and a four-species EcM plant community (*Castanopsis cuspidata* [Thunb.] Schottky, *Carpinus laxiflora* [Sieb. et Zucc.] Blume, *Pinus densiflora* Sieb. & Zucc., and *Quercus serrata* Murray) (Fig. 1a; Table S1 for information of ecological traits of individual species). The AM and EcM plant species used in this study occur sympatrically or parapatrically in secondary forests around Kyoto city, in which the experimental mesocosms of the model communities were constructed (see below).

For the conditioning phase described below (*Experimental design*), trees of the eight species (≤ 30 cm stem height), whose roots had been infected by naturally occurring mycorrhizal fungi, were acquired from native stands by a local nursery (Nakanishi-shiseien Co. Ltd.). Each tree sampling was placed in plastic pot until transported to glasshouse where we maintained all the tree samplings prior to the start of the experiment. Because conspecific trees possibly arrived at the glasshouse with different soil fungal community, we placed the “homogenizing” step that minimizes potential heterogeneity in fungal communities among conspecific trees before transplanting to the mesocosms for resident treatments. This increased the likelihood of our staring with a similar fungal community per host species, minimizing the posssibility that the fungal community differentiation occurred only among conspecific trees but not among heterospecific trees. To do this, the trees were grouped by plant-species and trees of each plant species (prospective conspecific resident trees in the experiment) were incubated for three months using different pots (60 cm × 40 cm × 20 cm) filled with “red-ball” soil (akadama) in a greenhouse prior to use in the experiment. During the pot treatment, each tree’s root system was covered by nylon mesh, allowing mycelia (but not fine roots) to spread across to neighboring conspecific trees. Trees were then transplanted to the mesocosms for resident treatments (with nylon mesh removed).

For the feedback phase described below (*Experimental design*), seeds were bought from the nursery and were individually sown in 9 cm^3^ pots filled with red-ball soil. To equalize the size of the seedlings of the eight species (5–10 cm stem height), the pots were maintained at 16°C for 3 months. We planted those uninoculated seedlings directly into mesocosms (instead of sowing seeds) in order to focus on treatment effects on the seedlings that survived, not on mortality before the development of mycorrhizal symbiosis. Three-month-old seedlings should be less susceptible than germinants to various environmental stresses (e.g., heat, frost, and desiccation) and were therefore suitable for our purpose.

### Experimental design

The common garden experiment was performed in an open field established at Kyoto University Botanical Garden, Kyoto city, Japan (35°07′49″N, 135°47′10″E). We designed a full factorial mesocosm experiment involving AM and EcM plant systems to assess and compare the effects of mycorrhizal type matching on the outcome of plant–soil feedbacks. A total of 36 replicate mesocosms were set up in four spatial blocks (each 4 m × 4 m), each block containing nine mesocosms. The treatment design included three resident levels (AM, EcM, and control [mimic trees]) × three seedling levels (AM, EcM, and control [no seedlings]) × four replicates in a randomized block design. In February 2012, we established the 36 mesocosms measuring 1.2 × 1.2 × 0.4 m^3^, filled with a standardized mixture of red-ball soil (70% of the volume), decomposed granite soil (20%), and litter soil (10%) (Fig.1b).

To set up the conditioning phase (Fig. 1c), three resident treatments were applied to mesocosms (Fig. 1a). For AM and EcM resident treatments, mycorrhizal trees were initially planted to establish whole fungal communities as conditioning live inoculum in the mesocosms (*Model system*). In doing so, we laid a grid system to create spatial heterogeneity in resident plant communities; each mesocosm was composed of 16 resident trees arrayed in 4 × 4 Latin square (Fig. 1a), with the constraint that all treatment plant species must be equally represented (i.e., 4 species × 4 individuals). The control-resident treatment comprised 16 handmade mimic trees (made of a pole, wire, and cheesecloth), potentially eliminating differences across treatments in the light conditions experienced by seedlings during the feedback phase. With this control treatment, we were able to focus on resident effects via alteration in nutrient competition and plant–soil feedbacks, rather than those via change in light availability.

At the feedback phase (i.e., starting 3 months after establishing resident communities; Fig. 1c), three seedling treatments were fully crossed with the resident treatments: introduction of uninfected AM seedling community (4 seedling species × 16 individuals), introduction of uninfected EcM seedling community (4 seedling species × 16 individuals), or no seedling addition. For seedling addition treatments, seedlings (one individual for each of 4 species) were planted directly under resident trees in each of the 16 grid cells within the individual mesocosm; in each grid cell, four seedlings were planted in the four inter-cardinal positions 5 cm from each of the centered resident trees, with seedling species’ positions assigned randomly in each grid cell (Fig. 1a). This configuration represented all possible species-pairwise resident–seedling interactions (within grid cells), thereby averaging resident–seedling interactions at the whole mesocosm scale. In the first week of the experiment, seedlings that died were replaced.

We made two modifications to the experimental design, because of low germination rates of two AM plant species (*C. camphora* and *P. jamasakura*). First, *C. camphora* was replaced by a surrogate AM species, *Celtis sinensis* Pers., for seedling treatment; hence this species occurred only as a seedling species, not as a resident species. Second, we transplanted *P. jamasakura* seedlings only to the second and third columns in mesocosms because we were not able to secure a sufficient number of the germinated seedlings.

### Growth survey and harvest

Over the two growing seasons, survival, height, stem width, number of leaves, and light intensity were followed for each seedling on a quarterly basis until the final census in September 2013. At the final census, leaf chlorophyll was measured for each seedling, using a SPAD502 Plus Chlorophyll Meter (Minolta Camera Co., Japan). At harvest, all resident trees and seedlings were carefully excavated with utmost care not to cut their roots, especially when resident and seedling roots intermingled. Each sample was thoroughly washed in a separate bucket filled with running tap water. For molecular analysis, we collected 2 cm pieces of about eight terminal roots from each washed sample (both of all the surviving seedlings and resident trees), and stored them in 99% ethanol at –20°C. All the raw samples (resident and seedling samples) were placed in plastic bags and stored in the refrigerator until further processing for scanning, weighing dry plant samples (e.g., leaf, root), and physiochemistry analysis. As a whole, we collected data for relative growth rate of seedling height, dry leaf and root weights (at harvest), total leaf area, seedling stem diameter at base, and leaf chlorophyll, but for illustrative purposes, we report the results of the analysis only for dry leaf and root weights.

### Fungal community structure

For each sample of resident trees and seedlings, the composition of root-associated fungi was examined using pyrosequencing. Each sample represented five randomly selected terminal roots (ca. 2 cm each) and was processed using the protocol detailed in the Supporting Information. Total DNA was extracted from each sample using the cetyltrimethylammonium bromide (CTAB) method. We amplified the entire internal transcribed spacer (ITS) region and the partial ribosomal large subunit (LSU) region using the fungus-specific high-coverage primer ITS1F_KYO2 and the universal primer LR3^35^. PCR was conducted with a temperature profile of 95°C for 10 min, followed by 20 cycles at 94°C for 20 s, 50°C for 30 s, and 72°C for 120 s, and a final extension at 72°C for 7 min using the buffer and polymerase system of Ampdirect Plus (Shimadzu). We subjected the PCR product from each root sample to a second PCR step that targeted the ITS2 region. The second PCR was conducted using the universal primer ITS3_KYO2 fused with 454 Adaptor A and sample-specific molecular ID; the reverse universal primer was LR_KYO1b fused with 454 Adaptor B. A buffer system of Taq DNA polymerase together with standard Taq buffer (New England BioLabs, Ipswich, MA, USA) was used with a temperature profile of 95°C for 1 min, followed by 40 cycles at 94°C for 20 s, 50°C for 30 s, and 72°C for 60 s, and a final extension at 72°C for 7 min. Sequencing was performed using a 454 GS Junior (Roche Diagnostics, Indianapolis, IN, USA).

We then performed bioinformatics analysis using Claident v0.2.2014.10.29^36^ as detailed in the Supporting Information. Briefly, fungal operational taxonomic units (OTUs) were defined at a cutoff sequence similarity of 97% and potentially chimeric OTUs were removed using UCHIME v4.2.40^37^. The taxonomic assignment was performed with the Query-centric auto-k-nearest neighbor method^36^ using the “nt” database downloaded from the NCBI ftp server (http://www.ncbi.nlm.nih.gov/Ftp/).

### Feedback effects on seedling growth

We calculated and compared feedback effects on seedlings across different treatments using data on the performance of the introduced seedlings in the mesocosms. To do this, the average seedling species performance was calculated in each mesocosm by all possible seedling and resident species combinations at the grid level, and the resultant population averages for different seedling species was used as a statistical unit for downstream analysis. For each of the four spatial blocks (of randomized block design, see *Experimental Design*), we used the population-average growth responses for different seedling species to formulate the feedback effects as follows: (P_consp_ – P_heterosp_)/P_control_, where P_consp_ represents seeding performance under conspecific resident species within the same grid (nearest neighbor), P_heterosp_ under heterospecific resident species, and P_control_ under no-resident control in the same spatial block. The feedback effect defined here is scaled to seedling growth in the no-resident control treatments (that is, artificial trees made of pole and cheesecloth), where seedlings grew in the absence of (i) soil nutrient competition with resident trees and (ii) mycelial inocula. For illustrative purposes, we analyzed data for final dry weights of leaf and root when calculating feedback effects, but the results were qualitatively similar when we used different growth components to represent feedbacks.

Using a linear mixed model (“lmer” function), feedback effects (i.e., P_consp_ – P_heterosp_)/P_control_) based on each component (a, leaf weight and b, root weight) were analyzed for each seedling species separately as a function of resident treatments as fixed effect, with spatial block (4 blocks) and resident species identities (4 species in each treatment) as random effects to account for species specificity in effects of resident species. We test the hypothesis that seedlings grown in matched resident environment perform better than seedlings grown with mismatched residents. Because *Celtis* was used only as seedling species but no as resident species due to modification of the experimental design, we could not estimate feedback effects for *Celtis*. To improve normality, feedback effects were square root transformed while retaining the sign. All statistical analyses were performed with R^38^, using the libraries “vegan”^39^ and “lme4” ^40^.

### Sharing of mycorrhizal fungi

Using molecular analysis of root-associated fungal communities on resident and seedling species (see Figs. S3-4), we examined whether seedling species had a greater chance of being colonized by root-associated fungi in matching resident treatments (AM resident/AM seedling and EcM resident/EcM seedling) than in mismatching treatments (AM resident/EcM seedling and EcM resident/AM seedling). Morisita-Horn similarity (1 – *C*_MH_) in fungal OTU composition was calculated between every possible seedling and resident species combination in individual mesocosms. Using a linear mixed model, the observed similarity was analyzed as a function of seedling and resident treatment types and their interaction term, with each plant species and spatial block being random effects. The mycorrhizal type matching effect was measured based on the resident type × seedling type interaction term.

### Spatial structuring of plant–fungus associations

To evaluate the tendency of seedlings to share OTUs with closer resident trees than with distant ones, we calculated compositional similarity using 1 – *C*_MH_ as a metric for every possible individual resident–seedling pair in each mesocosm. The metric was then quantified for its correlation with physical distances separating the pairs in individual mesocosms, using Spearman’s rank correlation ρ. A higher correlation indicates that closely located resident–seedling pairs shared a higher proportion of the fungal communities than did distantly located pairs. The correlation ρ was analyzed as a response variable, with seedling and resident treatment types and their interaction term as predictors. Because of the slightly different spatial design between AM and EcM seedling additions (see *Experimental design*), we examined differences in Spearman’s rank correlation coefficient ρ between resident types separately for each seedling type, using Welch’s *t*-test (instead of a linear model and two-way ANOVA).

## Acknowledgements

We thank Kyoto University Botanical Garden to support this work, and H Maki, H Chifuku, K Kitayama, M Mukai for help with chemistry analysis. Many students assisted with this experiment, and their help has been invaluable. We thank T Fukami, Y Onoda, K Po-Ju, T Miki for discussion, ALC Jousset, S Kefi for comments on this manuscript. This work is supported by Next Generation World-Leading Researchers of Cabinet Office, the Japanese Government (GS014), KAKENHI (26711026) and JST PRESTO (JPMJPR16Q6) to HT and JSPS Research Fellowship (13J02732) to KK.

## Authorship

KK, SY, and HT initiated and managed the project. KK and HT designed experiments. SY and HS performed DNA extraction, PCR and next-generation sequencing. AST performed bioinformatics analysis. KK analysed and interpreted all data. KK wrote the first draft with inputs from all the coauthors. All the authors contributed to revision of the manuscript.

## Data availability

The fungal ITS sequences of have been deposited in the databases under accession numbers BioProject PRJDB5467 and DDBJ DRA005499.

